# Population monitoring of snow leopards using camera trapping in Naryn State Reserve, Kyrgyzstan, between 2016 and 2019

**DOI:** 10.1101/2021.02.24.432722

**Authors:** Julie Rode, Claire Lambert, Lucile Marescot, Bastien Chaix, Julie Beesau, Suzanne Bastian, Joldoshbek Kyrbashev, Anne-Lise Cabanat

**Affiliations:** Objectif Sciences International NGO, OSI-Panthera program, Geneva, Switzerland; INRAE, Oniris, BIOEPAR, 44300, Nantes, France; Naryn State Nature Reserve, Kyrgyz Republic

**Keywords:** snow leopard, camera trapping, population dynamics, diachronic monitoring, citizen science, Kyrgyzstan

## Abstract

Four field seasons of snow leopard (*Panthera uncia*) camera trapping inside Naryn State Reserve, performed thanks to citizen science expeditions, allowed detecting a minimal population of five adults, caught every year with an equilibrated sex ratio (1.5:1), and reproduction: five cubs or subadults have been identified from three litters of two different females. Crossings were observed one to three times a year, in front of most camera traps, and several times a month in front of one of them. Overlap of adults’ territories was observed in front of several camera traps regardless of their sex. Significant snow leopard presence was detected in the buffer area and at Ulan, situated at the reserve border. To avoid poaching on this umbrella species and its preys, extending the more stringent protection measures of the core zone to both the Southern buffer area and land adjacent to Ulan is necessary.

## Introduction

Apex predators are important keystones of ecosystems’ viability ^1,2^. Hence, the long-term monitoring of their population census and dynamics in vulnerable landscapes is important to avoid the collapse of the trophic chain which damages ecosystems and has impacts on all living populations ^3,4^. Healthy Central Asia high mountain ecosystems have in general four top predators: bears (*Ursus arctos isabellinus*), wolves (*Canis lupus*), lynxes (*Lynx lynx isabellinus*) living at the lowest altitudes and snow leopards (*Panthera uncia*) living at the highest altitudes ^5^. Snow leopards are listed as Vulnerable (C1) by the IUCN ^6^ and their total world population is estimated at 2,710 to 3,386 mature individuals fragmented over twelve countries in Central Asia ^6^. In Kyrgyzstan, the total population was estimated at around 350-400 individuals (National Academy of Sciences of Kyrgyzstan: unpublished data)^5^. The Naryn State Reserve located in Kyrgyzstan is known for the protection of several ungulates and predators including snow leopards in the highest altitude ranges. A preliminary census giving five snow leopards per 200 km^2^ was performed in Naryn State Reserve ^5^. It was however not followed by a more precise estimation nor a long-term and systematic monitoring of this population. McCarthy and Mallone (2016) ^5^ recommended a more precise estimation and a long-term monitoring of this population, which led to this study. In addition, the Snow Leopard Survival Strategy written by members of the Snow Leopard Network emphasises the importance of providing accurate demographic trends ^7^.

As they are very elusive felids living in rocky and cliffy places ^5^ and as their range is covering very steep and not easily accessible ground, one of the best non-invasive methods to monitor their population is through camera trapping. This method is indeed widely used for recording and studying felids in general ^8,9^ and snow leopards in particular ^10–13^. Thanks to citizen science expeditions organized from 2016 to 2019, once to four times every summer, we were able to perform the first long-term monitoring of the snow leopard population in this region, in our sample area inside the reserve. This project, where volunteers help set up and retrieve pictures from camera traps and collect traces, is ongoing.

The main goal of this study is to put a first milestone on the assessement of the census and viability of the snow leopard population inside Naryn State Reserve. We also performed a first determination of their minimal territory inside our sample area, which can be shared with cubs; it shows overlapping between individuals. The detection and identification of individuals in Naryn State Reserve was based on a camera trap system deployed on a sample area inside the highest attitudinal range. The aim was to bring insights on their distribution and sustainability in the long-term. We were able to determine the yearly minimum population size, sex ratio, individual movements within our sample area as well as reproduction events with cubs recaptured by our camera traps over the years.

## Material and Methods

### Study area

The study area (**Figure 1**) is located inside Naryn region (Kyrgyzstan), in the eastern part of Naryn State Reserve created in 1983. The altitude inside the reserve varies between 2200 and 4500 meters ^14^. Most of the study area is located in the core zone of the reserve while a small area, including Kok-ozon valley, is situated in the buffer zone. The North-Eastern part is bounded by the Naryn and Ulan rivers, while the ridges of the Naryn-too mountain range occupy the Southern limit. To the West, the study area is bounded by the Chon-taldo valley. Habitats are composed of spruce forests up to 3000 meters on the Northern slopes, while the rest of the landscape is occupied by mountain pastures, rocky areas and glaciers ^14^. No human activities are authorized except accredited scientific activities. Extensive pastoralism is allowed in the buffer zone, but not in the core area. On the surroundings of the reserve, extensive pastoralism, tourism and sport hunting are present. The entire reserve covers an area of 1056 km^2^ inside which the study area occupies approximately 260 km^2^.

**Figure 1.**
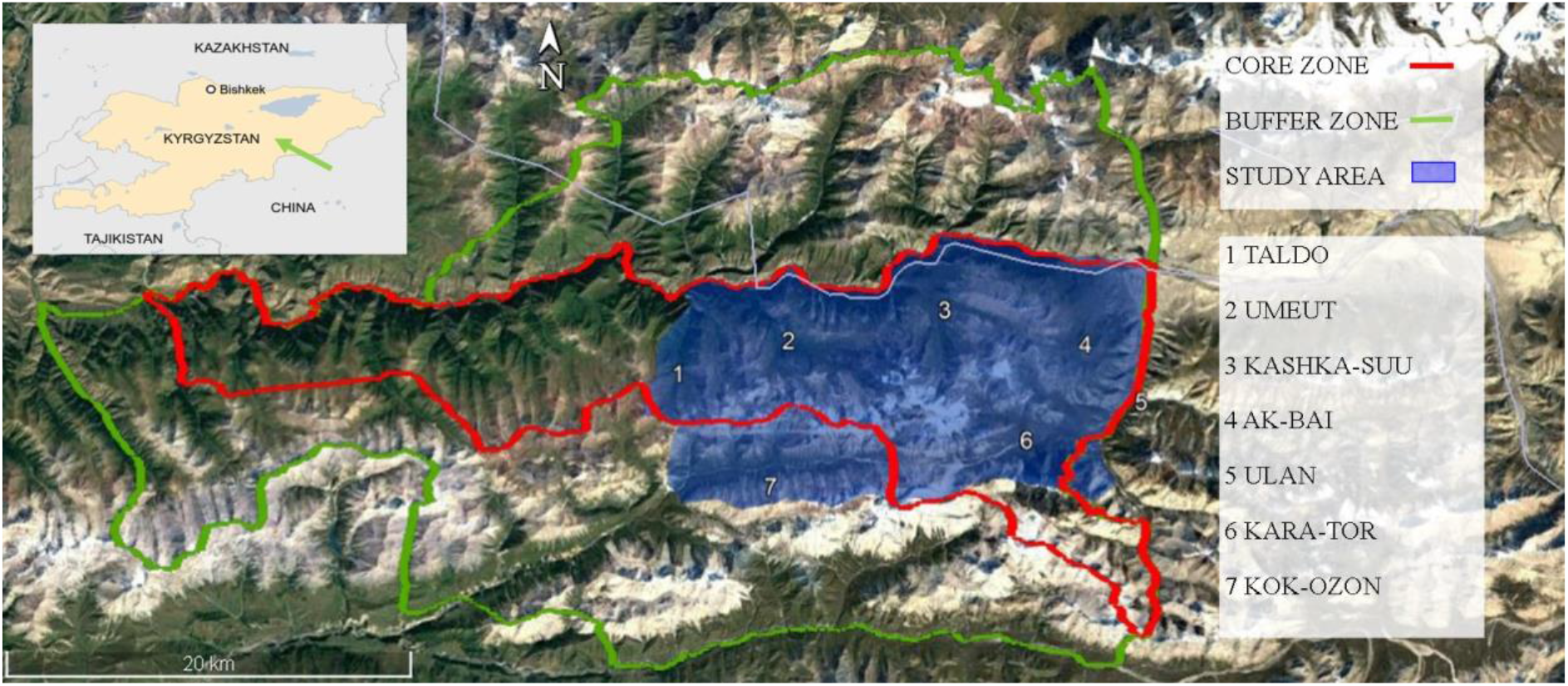
Map of the study area in and around Naryn State Reserve

### Camera trap on-field dispositive

Camera traps were set up and pictures were retrieved during citizen science expeditions, under the supervision of scientists and reserve rangers. We used Bushnell Essential (E2 and E3) camera traps with infrared sensor and low glow LED flash. They were equipped with SD cards of at least 16 gigabits and lithium batteries. They stayed in position all year long.

The camera traps were installed in permanent stations, within about five kilometers from one another ^11^, in favorable places where we found snow leopard signs of presence (pugmark, scat, scrape, urine spray or hairs), on narrow passes, along cliffs, facing large rocks, etc. ^10^. Each camera trap is fixed on a rock with a strap at around fifty centimeters to one meter from the ground and at a minimum of two meters from the expected snow leopard crossing. When possible, orientation is chosen with a horizontal angle of about 45° to the estimated snow leopard crossing location.

They are set up on camera mode, with bursts of three pictures, a 0.3 second trigger between each picture and one second between each burst, and without time lapse or field scan. They register fauna on the spot, all year long except if unintentional triggers saturate the SD card and exhaust the batteries or, if the camera is damaged by fauna. The camera traps are checked at least once every summer, sometimes more often: SD cards are replaced, new batteries are installed and if necessary, the position and orientation are adjusted on the same site or to a new one.

The study area was expanded between 2016 and 2017. Three camera traps were set up in 2016. In 2017, we added new camera traps bringing them to a total of ten. In 2018 and 2019, nine camera traps were running simultaneously. Activity locations and dates are available in **Table S1** in the supplementary materials.

### Manual Individual Identification

As software identification is not efficient enough yet to properly differentiate individual snow leopards ^15^, we performed manual identification based on the unique patterns of their fur rosettes ^10,11,13,16,17^ (**Figure 2**). Three distinct readers made an independent examination of all 1984 retrieved snow leopard pictures following the identification principles stated for bobcats ^18^ and already used for snow leopards ^10^.

**Figure 2.**
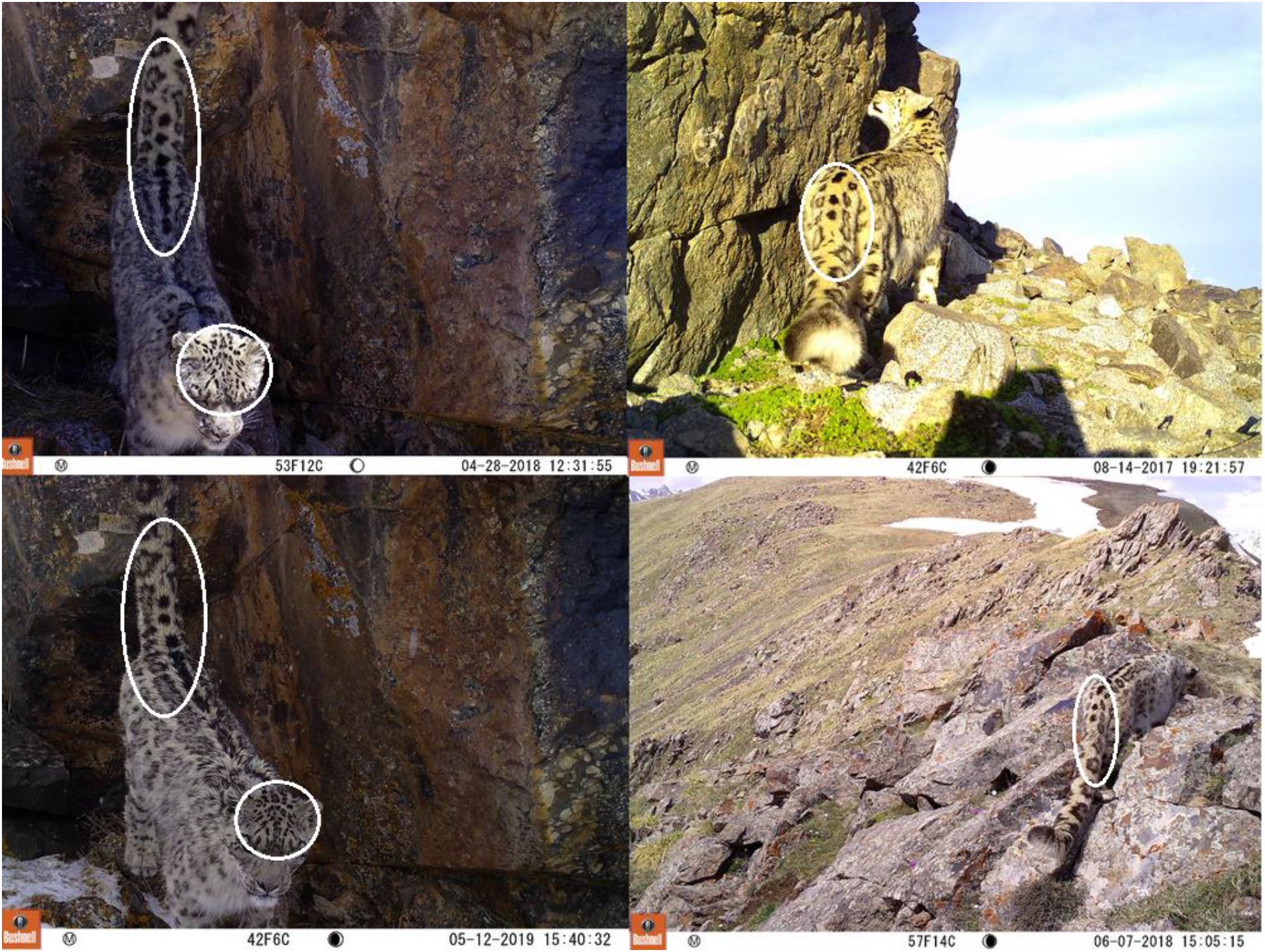
Pictures of Naguima from OSI-Panthera camera traps with highlighted areas with similar rosettes

The results of each reader’s analysis were put in common to obtain the final identification. In case of disagreement on the identification of an individual in a picture, the picture was discarded. For each individual a picture library was created, a name was given, the recording location and time, the sex when determined, and the presence of other individuals were noted. This allowed for a growing number of pictures of each individual over the years which in turn made possible the a posteriori identification of individuals. Sex determination was made only when we could observe the presence or absence of genitals on pictures ^19^ or in case of a mother with cubs, as they stay together during one to two years of breeding ^20,21^.

We consider one “crossing” or “event” when a snow leopard, a couple or a mother and its litter, is pictured by a camera trap. We consider that two different events have occurred, when more than 15 minutes have passed between two pictures of a same individual or group.

### Software analysis

Data concerning each snow leopard, namely time, location of observation (camera trap number, position, GPS coordinates), sex and if accompanied, are compiled into a single database. R software ^22^ was used for extracting data from the database and performing statistics, visual representations and mapping using *ggplot* ^23^ and *tidyverse* ^24^ packages.

We calculated the Minimum Convex Polygon (MCP) for individuals detected at least in three different camera trap stations. We used the function “mcp” from the package *adehabitatHR* ^25^ which draws the smallest area used by an individual defined so that all the detection points within the polygon have interior angles inferior to 180 degrees. MCP is a common estimator of animal home range ^26^, but we will use it in this study to determine the minimal territory of individuals inside our sample area. It will describe the potential areas used by animals identified by cameras in our sample area, acknowledging that some areas used by individuals may have been overlooked (individuals not detected but yet present) or that other areas which are actually never used may have been wrongly included.

## Results

Each year we captured snow leopards on most of our camera traps. The Ortho Taldo camera trap was the only one which never captured snow leopards. Among the 172 snow leopard crossings observed, we were able to precisely identify individuals on 156 crossings. Identification wasn’t possible when only fur appeared on the picture or when individuals were covered by snow. **Table 1** shows which snow leopards were captured each year. In 2019, the minimal number of individuals in our study area was five adults and three subadults. All adults had been recaptured during previous years (**Table 1**). Among them we identified three females and two males. **Figure 3** displays the minimal number of adults, cubs (less than one year) and subadults (more than one year still with their mother) that have been captured between 2016 and 2019. Adults together seem to have been caught only once. There were three none-identified adults or, at least one adult with two subadults, covered by snow at Ortho Kara-tor in 2017.

**Table 1:** Capture and recapture of each identified individual along the years of the study with sex if determined

|  | Individual | Aika | Ayana | Fatima | Ilguiz | Malie | Merim | Naguima | 3 cubs<br>Aika | 1 subadult<br>Ilguiz |
| --- | --- | --- | --- | --- | --- | --- | --- | --- | --- | --- |
|  | Sex | Female | Female | / | Female | Male | / | Male | / | / |
| **Capture /**<br>**Recapture** | 2016 | x |  | x | x | x | x | x |  |  |
|  | 2017 |  | x |  | x | x |  | x | x |  |
|  | 2018 | x | x |  | x | x |  | x | x | x |
|  | 2019 | x | x |  | x | x |  | x | x |  |

**Figure 3.**
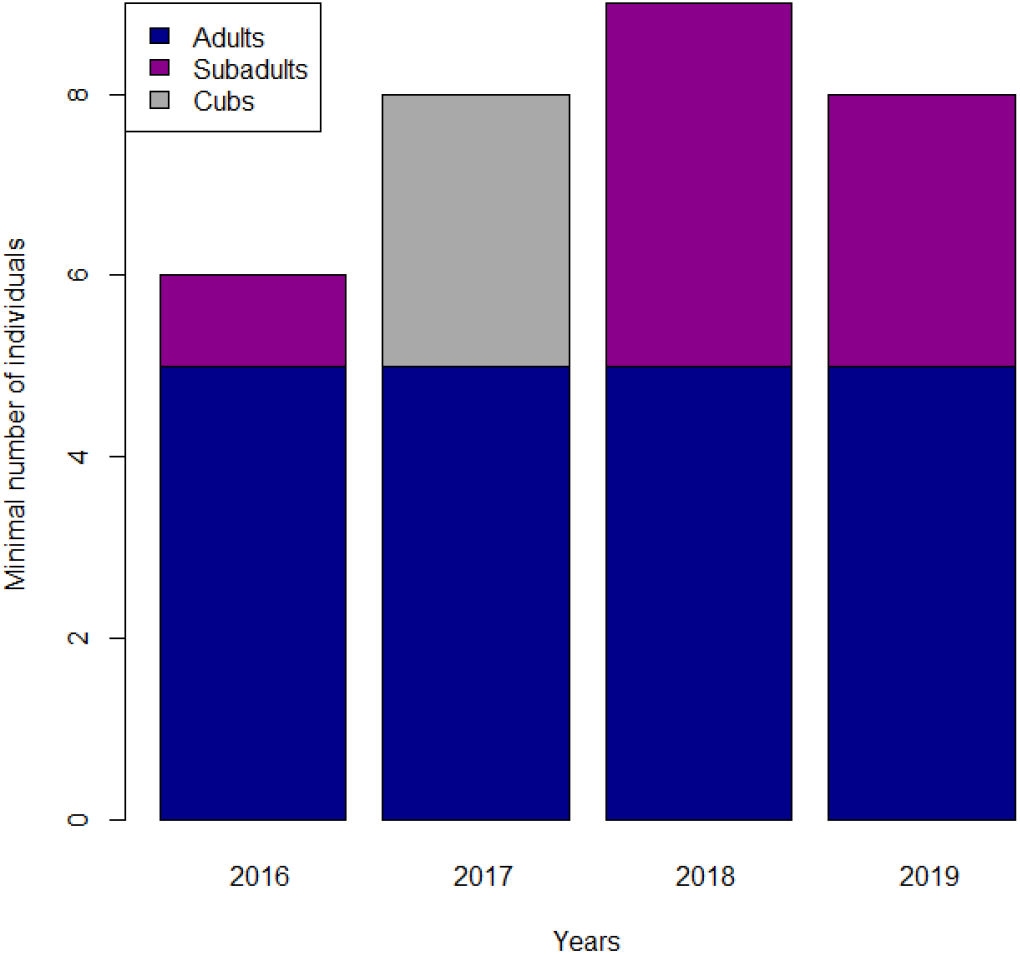
Minimal number of captured Snow Leopard along the years of the study divided between adults (blue), subadults (purple) and cubs (grey). Aika was taken into account in 2017 even if not pictured as she was present before and after in our camera traps and pictured by KWRCPSUNSNR camera traps

Two of the three identified females have been captured with cubs during the four years of study. The individual named Aika was observed in 2016 with a 1 to 1.5 years old subadult Merim and from 2017 to 2019 with another litter of three cubs. Based on their morphology, the birth of these three cubs occurred during spring 2017. The individual Ilguiz was observed with a subadult in Ulan Kara-tor in May 2018.

Yearly trajectories taken by the individuals recaptured across camera traps are shown on **Figure 4**: three individuals travel extensively through the reserve, Aika (female, detected by five camera traps), Ayana (female, by six camera traps) and Naguima (male, by eight camera traps); others are seen on a smaller portion of the map. One individual, Ilguiz (female), was detected every year at a single camera trap location. Ilguiz’s cub was also detected elsewhere and was likely following its mother.

**Figure 4.**
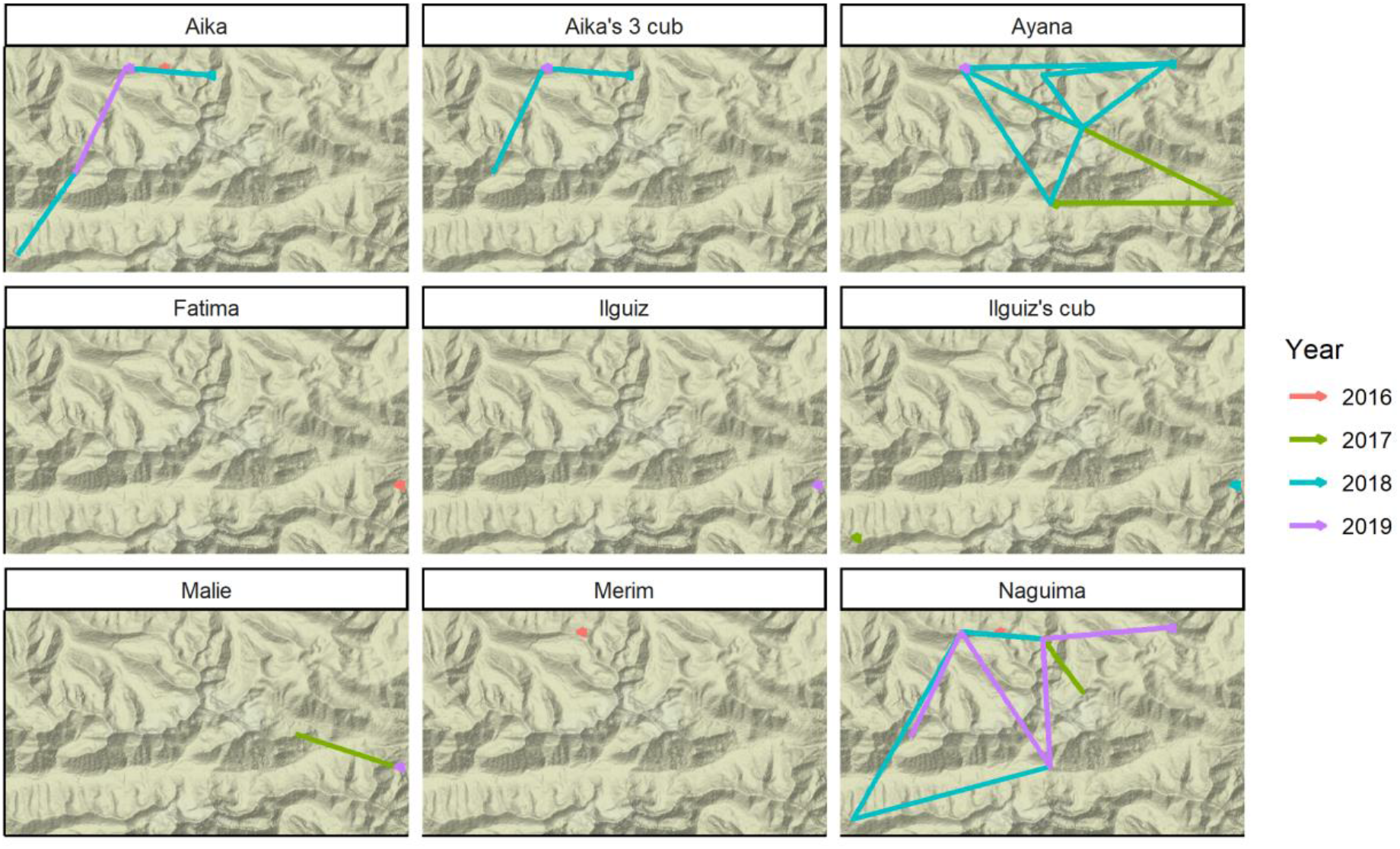
Individual movements between the different camera traps by year

We could estimate the minimal territory inside our sample area of only three individuals using the MCP function. The male Naguima has the largest MCP, followed by the female. The female Aika seems to use a smaller proportion of the study area. The three polygons were partially overlapping (**Figure 5**). Five individuals were captured by the camera trap at Umeut, with a total record of 95 crossing events over the entire study period. The camera trap at Kashka suu glacier has the least number of crossings: two individuals (the male Naguima and female Ayana) were identified in three crossing events (**Table 2**).

**Table 2.** Number of crossings and unique individuals captured by each camera trap location.

| Location | Number of snow<br>leopard's crossings | Number of<br>identified<br>individuals |
| --- | --- | --- |
| Kok-ozon - Ailuteur | 5 | 4 |
| Umeut | 4 | 3 |
| Akuluk 2 | 9 | 4 |
| Umeut - ridgeline | 95 | 5 |
| Ulan Dungeureumeu/Kara-tor | 21 | 5 |
| Kok-ozon pass | 6 | 2 |
| Kashka suu glacier | 3 | 2 |
| Ak bai | 6 | 2 |
| Kashka suu West | 23 | 4 |

**Figure 5.**
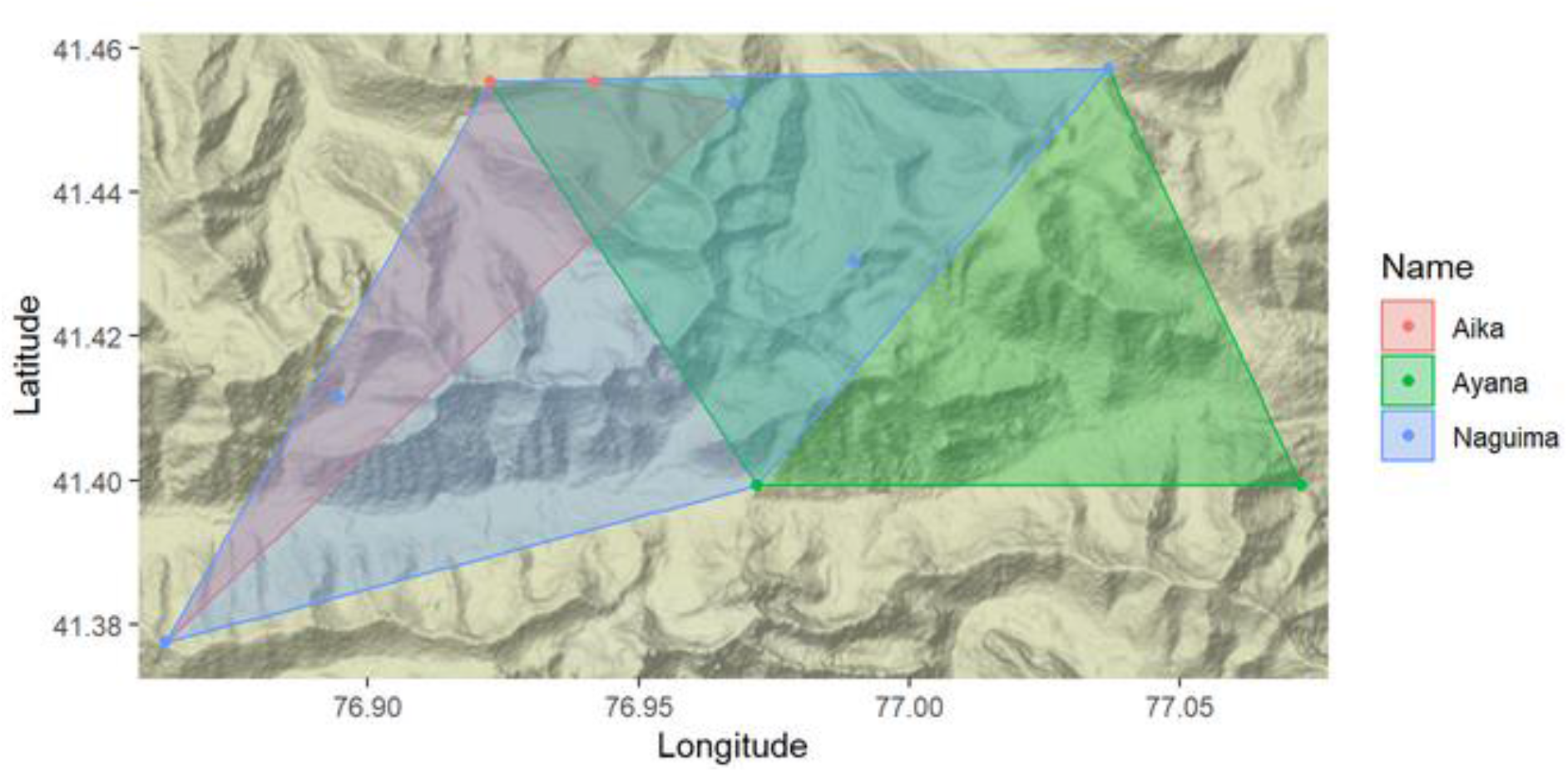
Minimal territories in our study area of Aika, Ayana and Naguima determined by using the Minimum Convex Polygon method and the camera-crossings

Frequent crossings were observed in front of several camera traps. Naguima crosses one to three times along the year in front of each camera trap at the exception of Umeut. At Umeut it crosses the camera trap every month, sometimes three times a month with an interval of around 10-20 days (a lack of data in the Umeut area for the first half of 2017 due to a camera trap problem has to be noted). The same trend is observed for Ayana and Aika which cross in front of most of the camera traps one to three times a year with a highest frequency in the Umeut area. Ayana crossed two to three times a month from August to October 2018 and from January to March 2019. Aika was pictured every month since August 2018, one to three times a month, except in March 2018, November 2018 and March 2019. All three of them were observed alone, or with cubs in Aika’s case. At Umeut, however, Aika and Naguima were pictured twice at 11 hours (07 May 2018) and at 20 minutes (25 May 2018) intervals.

In **Figure S1** of the Supplementary materials, we summarized the percentage of crossings by hour. The maximal activity for snow leopards took place between 6PM and 8PM (around 30% of observations) and the minimal activity between 11AM and 4PM (around 10 % during these five hours). This result has a sampling bias and will be improved by using appropriate methods in a dedicated spatially-explicit capture-recapture model study ^27^.

## Discussion

This survey gives an insight into the snow leopard population census and dynamics inside Naryn State Reserve. From 2016 to 2019, we observed five individually identified adults in our sample area of around 260 km^2^. In another reserve in Kyrgyzstan, Sarychat-Ertash, the population inside a sample area of 1,341 km^2^ was estimated at 18 individuals by genetic analysis of scats ^27^ and in 2014, 15 individuals were counted by camera trapping ^5^. However, at this time we can’t compare our observed number of individuals in terms of population densities to these numbers as our sample area is too small to compute it accurately.

Among the five adults, we identified three females and two males i.e. a sex ratio of (1.5:1). However, given the small census and the high variability of this ratio ^11^ this evolution should be monitored over a longer period of time, to ensure the population remains sustainable in the long term. We caught reproduction events during the four years of study (**Table 1**). One of Aika’s litters was observed during two years (spring 2017 to spring 2019) with its mother. This falls within the usual range of breeding period during which juveniles stay with their mothers ^21,28^. Aika strongly contributes to the reproduction of the population. We have already registered three litters for her: two subadults in 2014 (data from the Kaiberen Wildlife Research and Conservation Project of the Shinshu University and Naryn State Nature Reserve (KWRCPSUNSNR)), one subadult (Merim) in 2016 and three cubs born in 2017. In captivity, litters are usually composed by one to five cubs, with a mean of 2.1 cubs ^29,30^, which is in line with our observations. In the wild, in Mongolia the survival probability before two years of age has been estimated at 0.83 (+/−0.15) ^11^. In this study, the survival rate is good as in five years, Aika brought up six cubs to adulthood and raised up to three of them at the same time. However, we lost track of Merim and of Ilguiz’ subadult, hence, we don’t know if they reached adulthood. They could still be there, either observed but not identified (blurred pictures or pictures showing the wrong profiles for instance) or unobserved. They could also have left the area to settle far from their parents’ territory ^28^. The sexual maturity of snow leopards is reached at around two to three years old ^31–33^ and as Aika was caught with subadults for the first time in 2014, we can estimate that in 2019 Aika was at least eight years old. Another female, Ayana was photographed during summer 2016 by camera traps from the KWRCPSUNSNR. She then had the size of a subadult/adult and she was with another non-identified adult female. Hence, Ayana was at least five years old in 2019.

We were able to follow Aika’s litter since 2017 and they are now old enough to have similar fur to the one they will have as adults ^34^. Their monitoring in the following years will be easier if they do not leave the area. The knowledge on this new generation will allow to better estimate the existence of surrounding individuals and the possibility of new snow leopards entering the study area going against inbreeding possibilities. Indeed, as we are not aware of the snow leopards’ demography in the neighbourhood of our sample area and as we don’t have any mating pictures or details, we can’t conclude on that matter. Further genetic analyses in process on faeces samples could help shedding light on this issue.

We recaptured the same individuals every year so our saturation curve for adults is flat and in addition we witnessed reproduction events (**Figure 3**). We will however need a more detailed analyse to estimate the long-term stability of this population. We probably caught most of the study area’s population which validates our camera trap system. This system is widely used in other camera trap studies on snow leopards ^11^. As recommended by the Snow Leopard Network future years will see an expansion of our study area to reach 500 km^2^, with 30 to 40 camera traps.

Aika, Naguima and Ayana appear on a large number of camera traps throughout the years meaning that their territories are fixed and, at least partially, inside our sample area (**Figure 4**). The other adults, Ilguiz and Malie, have territories that are tangent to our sample area as they are seen only in one or two camera traps situated at the edge of the studied area.

Pictures from camera traps in different locations showed several individuals supporting the idea that adult snow leopards have overlapping territories, regardless of their sex ^35,36^. Snow leopards are rarely observed with other adults except when mating and for mothers with cubs during the breeding period ^28,35,36^. This is supported by pictures of Aika and Ilguiz with their respective cubs and by pictures from a KWRCPSUNSNR camera trap where a male and a female, both subadults, were photographed together in the Kashka suu area in June 2014. However, exceptions to those empirical observations were witnessed ^35,36^ and we didn’t observe more of Aika’s offsprings in the following months. Some of our individuals cross in front of the same camera traps several times a year. This suggests that they regularly go to the same places and use what is called travel routes for hunting, controlling and going through their territories ^5^. Some places are however more frequently used, like Umeut for Naguima, Aika and Ayana. The Kashka suu West camera trap also showed a high frequency of crossings. Both spots, particularly Umeut, seem to be strategic areas where the territories of three snow leopards (male and female) overlap and where females are present with their litters. It could be explained by the high presence of snow leopards’ preys in these areas. We observed for instance large herds of ibexes in the Kahska suu area and ibexes and argalis in Kok-ozon, Kara-tor and Ulan.

In wildlife surveys based on GPS telemetry, MCPs are commonly used to estimate animals’ home range ^37^, but in this study it was only used for displaying the minimal territory of each individual inside our sample area. However, MCP is subject to significant bias. It can include areas not used by animals and thereby overestimate home range sizes. It can also overlook areas where the animal is present but not observed ^38^. Those biases are more likely to occur when MCP is calculated from camera-trap studies ^39^. Therefore, we remained cautious on the interpretation of the results on individuals’ MCPs (**Figure 5**) and only observed trajectories across camera traps. For this reason, we deliberately chose to plot the MCPs without displaying metric estimations. Indeed, an individual observed at only one camera trap may have a larger minimal territory inside our sample area than for instance Naguima, which showed the largest MCP and greatest number of crossings among camera traps. Indeed, for some behavioral or environmental reasons, an individual can be observed in only one camera trap but missed being detected in other camera traps, despite occupying the surrounding areas. Thereby, individual heterogeneity in the detection may bias estimates. For instance, we found subadults in some camera traps but never detected the mother, which was likely near her dependent cubs. This highlights the importance of accounting for the detection probability, to make accurate estimations of minimal territories, minimal territories overlap between individuals and in general animals’ movements across the study area ^40^. Maintaining our current camera-trap monitoring and extending it on the long-term will allow us to collect more data and further estimate population density and dynamics using a spatially-explicit capture-recapture model for open populations ^41^.

No snow leopard was photographed by the Ortho Taldo camera trap but rangers have previously retrieved pictures from this valley. The South West of the study area extends into the buffer zone and shows four crossings by four individuals (**Table 2**) meaning that we also have snow leopards in the Southern buffer area. Ulan, situated at the edge of the reserve, also shows crossings from four different individuals. This place is adjacent to a hunting concession. In fact, Naryn State Reserve was initially created to protect the spruce forest habitat and the Tian Shan wapiti (*Cervus canadensis songaricus*) ^14^. Snow leopard protection came after; hence borders are not adapted to this species. Shepherds grazing inside the buffer area and ungulates hunting at the borders of the reserve can put pressure on the snow leopardss’ preys. It also provides more opportunities for poaching both snow leopards and their preys. Consequently, to increase the protection of the species and its preys we strongly weigh in for the extension of the core zone to include at least the Southern buffer area and a large portion of the land adjacent to Ulan.

In addition, places where a lot of crossings by many individuals were registered raise the question of the strategic location of such places and of territories overlapping. Monitoring with genetic samples could allow for a broader repartition of collected faeces’ samples, give a better idea of the real territories of each individual and provide some insights on these “hot spots”.

Looking at the time of day when the pictures were taken shows that this species is animated during crepuscule and quietter through the day which is coherent with other observations ^36,42^.

In addition to the usual threats coming from poaching and the reducing of their preys, inbreeding, if validated as a threat to this species, could also be a menace if there is no migration. Further genetic analyses in process on collected faeces’ samples and a significant extension of the study area of camera trapping would allow to conduct relatedness analysis, make a spatially-explicit capture-recapture model and gain more insights on minimal territory overlapping between males and females, reproduction success and generally, on the dynamics and distribution of the whole population^43^. Finally, camera trapping combined with citizen science seems to be relevant to perform a long-term non-invasive monitoring at low cost. This kind of study relies on volunteers who can also participate in other aspects of the data collection: scat sampling, ungulate counting, etc. It is consequently not dependent on external fundings and thus can be sustainable. It is also a way to develop responsible tourism and environmental education in snow leopard range countries, which will strengthen conservation efforts.

## Acknowledgments

We would like to acknowledge Naryn State Reserve director Mirtemir Turdaliev Mukambetovitch, vice-director Joldoshbek Kirbachiev and rangers for supporting us during our studies in the reserve, Maksatbek Anarbaev from the Kaiberen Wildlife Research and Conservation Project of the Shinshu University and Naryn State Nature Reserve for sharing some of their camera trap data from 2016 to 2019, our local partners, all the OSI-Panthera expedition volunteers and staff members of *Objectif Sciences International* NGO. We would also like to acknowledge Fréderic Salgues from Piege Photographique for providing us the camera traps at reduced prices, Rosina Karavia for spell checking and Koustubh Sharma for his helpful insights.

## Author’s contributions

J.R., B.C., A.-L. C. designed the study. J.R., C.L., B.C., J.B., J.K. and A.-L. C. participated in the field camera trapping. J.R., C.L., B.C. replaced in 2019 by J.B. perfomed the manual identification of individuals. J.R, C.L., L.M. and A.-L. C. analyzed the data. J.R. and A.-L. C. wrote the manuscrit. All authors revised the manuscript and approved the final version.

## Ethics approval and consent to participate

An authorization to set camera traps was given by the head of Naryn State Reserve and an autorization to mention some camera trap results from the Kaiberen Wildlife Research and Conservation Project of the Shinshu University and Naryn State Nature Reserve was given by Maksatbek Anarbaev.

In the OSI participation conditions (http://www.vacances-scientifiques.com/Conditions-de-Participation.html), it is stated that data gathered by participants during these expeditions will be used for scientific purposes.

