## Supplementary materials for "Population monitoring of snow leopards using camera trapping in Naryn State Reserve, Kyrgyzstan, between 2016 and 2019"

**Table S1**: Camera trap locations and main habitat along the study. For each location dates of activity are highlighted in dark grey and dates of inactivity due to damages are highlighted in light grey.

| ***Location*** | ***Habitat*** | **2016** | | **2017** | | | | **2018** | | | | **2019** | | | |
|---|---|---|---|---|---|---|---|---|---|---|---|---|---|---|---|
| | | ***Summer*** | ***Autumn*** | ***Winter*** | ***Spring*** | ***Summer*** | ***Autumn*** | ***Winter*** | ***Spring*** | ***Summer*** | ***Autumn*** | ***Winter*** | ***Spring*** | ***Summer*** | ***Autumn*** |
| Kok-ozon - Aïluteur | Crest | | | | | | | | | | | | | | |
| Umeut | Crest | | | | | | | | | | | | | | |
| Umeut - ridgeline | Crest | | | | | | | | | | | | | | |
| Akuluk 2 | Valley | | | | | | | | | | | | | | |
| Ulan Dungeureumeu | Valley | | | | | | | | | | | | | | |
| Ulan Dungeureumeu /Kara-tor | Valley | | | | | | | | | | | | | | |
| Ortho Kara-tor | Crest | | | | | | | | | | | | | | |
| Kok-ozon pass | Pass | | | | | | | | | | | | | | |
| Ortho taldo | Crest | | | | | | | | | | | | | | |
| Kashka suu glacier | Crest | | | | | | | | | | | | | | |
| Kashka suu West | Crest | | | | | | | | | | | | | | |
| Ak baï | Pass | | | | | | | | | | | | | | |

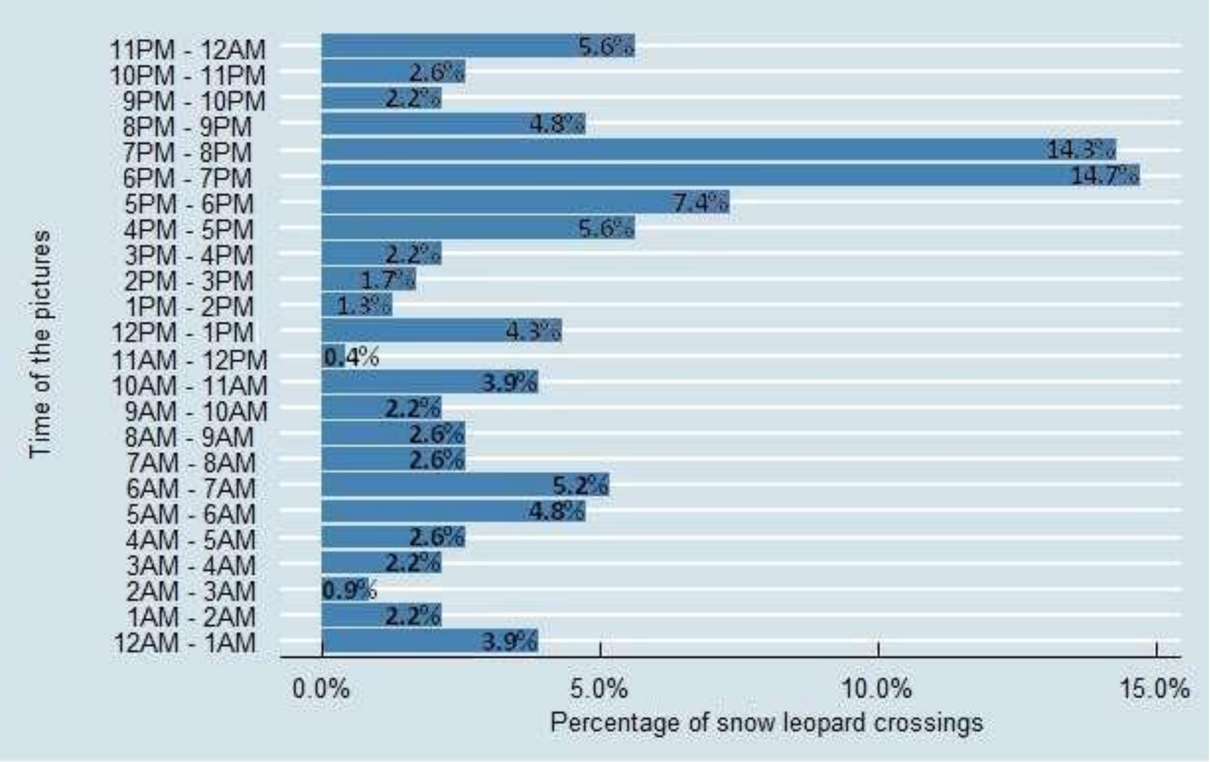


**Figure S1**- Summary of crossings by hour of day, from a total of 172 events.
